# Spherical harmonics texture extraction for versatile analysis of biological objects

**DOI:** 10.1101/2024.07.25.605050

**Authors:** Oane Gros, Josiah B. Passmore, Noa Borst, Dominik Kutra, Wilco Nijenhuis, Timothy Fuqua, Lukas C. Kapitein, Justin Crocker, Anna Kreshuk, Simone Köhler

**Affiliations:** European Molecular Biology Laboratory, Cell Biology and Biophysics Unit, Heidelberg, Germany; Cell Biology, Neurobiology and Biophysics, Department of Biology, Faculty of Science, Utrecht University, Utrecht, The Netherlands; Centre for Living Technologies, Alliance TU/e, WUR, UU, UMC Utrecht, Utrecht, The Netherlands; European Molecular Biology Laboratory, Developmental Biology Unit, Heidelberg, Germany

## Abstract

The characterization of phenotypes in cells or organisms from microscopy data largely depends on differences in the spatial distribution of image intensity. Multiple methods exist for quantifying the intensity distribution - or image texture - across objects in natural images. However, many of these texture extraction methods do not directly adapt to 3D microscopy data. Here, we present *Spherical Texture* extraction, which measures the variance in intensity per angular wavelength by calculating the Spherical Harmonics or Fourier power spectrum of a spherical or circular projection of the angular mean intensity of the object. This method provides a 20-value characterization that quantifies the scale of features in the spherical projection of the intensity distribution, giving a different signal if the intensity is, for example, clustered in parts of the volume or spread across the entire volume. We apply this method to different systems and demonstrate its ability to describe various biological problems through feature extraction. The *Spherical Texture* extraction characterizes biologically defined gene expression patterns in *Drosophila melanogaster* embryos, giving a quantitative read-out for pattern formation. Our method can also quantify morphological differences in *Caenorhabditis elegans* germline nuclei, which lack a predefined pattern. We show that the classification of germline nuclei using their *Spherical Texture* outperforms a convolutional neural net when training data is limited. Additionally, we use a similar pipeline on 2D cell migration data to extract polarization direction, quantifying the alignment of fluorescent markers to the migration direction. We implemented the *Spherical Texture* method as a plugin in *ilastik*, making it easy to install and apply to any segmented 3D or 2D dataset. Additionally, this technique can also easily be applied through a Python package to provide extra feature extraction for any object classification pipeline or downstream analysis.

**Author summary:** We introduce a novel method to extract quantitative data from microscopy images by precisely measuring the distribution of intensities within objects in both 3D or 2D. This method is easily accessible through the object classification workflow of *ilastik*, provided the original image is segmented into separate objects. The method is specifically designed to analyze mostly convex objects, focusing on the variation in fluorescence intensity caused by differences in their shapes or patterns.

We demonstrate the versatility of our method by applying it to very different biological samples. Specifically, we showcase its effectiveness in quantifying the patterning in *D. melanogaster* embryos, in classifying the nuclei in *C. elegans* germlines, and in extracting polarization information from individual migratory cells. Through these examples, we illustrate that our technique can be employed across different biological scales. Furthermore, we highlight the multiple ways in which the data generated by our method can be used, including quantifying the strength of a specific pattern, employing machine learning to classify diverse morphologies, or extracting directionality or polarization from fluorescence intensity.

## Introduction

Patterns are widespread in nature and can be observed across scales from subcellular to tissue and organism level. The complex interactions and mechanisms that underlie pattern formation processes are a topic of great interest in various fields (Rombouts et al., 2023). On a tissue- or cellular-scale, pattern formation is captured through 2D or 3D microscopy. Analyzing patterns in such images requires specific image analysis tools. One such class of analysis tools is texture extraction tools that describe the pattern, or generally, the morphology of biological systems, in microscopy images as a texture: the variation in signal intensity across an image (Armi & Fekri-Ershad, 2019). A number of different methods to extract texture information from images currently exist (Armi & Fekri-Ershad, 2019; Humeau-Heurtier, 2019). However, many of the methods rely on 2D natural images and cannot readily be applied to 3D biological microscopy data. Additionally, with the recent rise in accessible 3D microscopy segmentation methods (Berg et al., 2019; Stringer et al., 2021; Weigert et al., 2020), the number of applications in biology for accessible texture extraction from 3D data has risen. This need is shown by the different solutions using frequency space quantification for cell-cortical intensity (Mazloom-Farsibaf et al., 2023) and cell shape (van Bavel et al., 2023).

For many biological systems, understanding and quantifying 3D morphology throughout the object is a prerequisite to gaining new insight into different processes. For example, well-described developmental pattern formation, such as those arising during *Drosophila melanogaster* embryogenesis, are regulated by complex gene regulatory networks. The gene *shavenbaby* (*svb*) produces a striped expression pattern in the epidermis of the embryo, which later induces the formation of trichomes (Payre et al., 1999). Molecular changes in the upstream enhancers of *svb* have been shown to perturb the expression pattern, which can shape morphological evolution (Frankel et al., 2011) (Fig. 1A). To understand the phenotypic effects caused by sequence variations in regulatory elements, it is essential to analyze deviations from the typical wild-type expression pattern. Image texture can also provide biological information in systems where the pattern is not predefined but an emergent result of mechanical factors. A prime example here is the different chromatin morphologies that characterize the different substages of meiotic prophase I. These distinct substages, along with their corresponding DNA morphologies, are easily identified in the germline of the nematode *C. elegans* (Fig. 1B). As these varied morphologies directly correspond to the underlying molecular processes, any discrepancies in the spatial distribution of these morphologies can serve as indicators for detecting defects in meiotic timing (Hillers et al., 2017).

**Figure 1.**
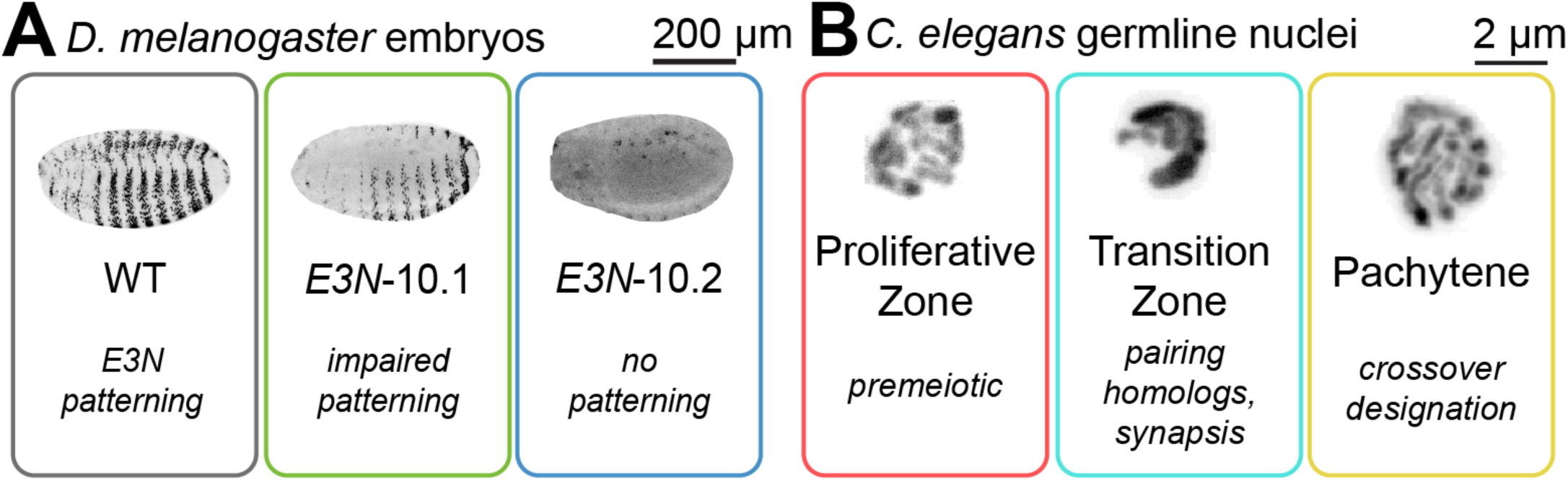
Patterns and image texture reflect biological spatial variability. **A)** Expression patterns of a Lac-Z reporter controlled by three different variants of the *E3N* enhancer: wild-type E3N, showing the expected striped ventral ‘*shavenbaby*’ phenotype patterning, the *E3N* mutant 10.1 with 10 mutations in *E3N*, with impaired patterning, and the *E3N* mutant 10.2 with 10 other mutations in *E3N* that lacks the patterning. These phenotypes reflect how random mutations disrupt the regulatory capacity of the *E3N* enhancer. **B)** *C. elegans* germline nuclei change DNA morphology during meiotic prophase I. The cells remain in the proliferative zone, showing small DAPI patches until they complete meiotic S-phase. In the transition zone, the chromatin is clustered as homologous chromosomes pair and co-align through synapsis. They then separate into strands of paired homologs in pachytene, as they designate the locations of crossovers, which are recombination events between maternal and paternal DNA.

Although their overall appearance is highly distinct, both the morphology of a fly embryo and of nematode meiotic nuclei can be described as a radial variance in fluorescence texture from their center of mass, allowing for robust quantitative analysis.

In this paper, we present a texture extraction tool for segmented microscopy data through frequency analysis of radial variation of image intensity. Our analysis assumes that the analyzed objects are mostly convex, which is true for many biological systems and has been a basic assumption for other algorithms (Weigert et al., 2020). We show that our texture extraction method can detect patterns in *D. melanogaster* embryos and distinguish different morphologies in *C. elegans* nuclei. Simple machine learning models trained with this feature perform as well in the classification of *C. elegans* germline nuclei as convolutional neural network models, while being faster to train. We also include a 2D implementation that allows us to quantify the actin leading edge of cultured cells and gives options for subsequent signal analysis for directionality mapping.

To make this method accessible, the method is implemented as a plugin for the user-friendly graphical software *ilastik* (Berg et al., 2019). Our implementation allows users to combine the *Spherical Texture* feature with other image features and quickly assemble a simple Random Forest classifier, which can be interactively trained within the program.

## Results

### Spherical Texture Method

In *C. elegans,* the condensation and organization of chromatin in the nucleus changes throughout meiosis. The nucleus shown in Fig. 2A is in pachytene, where pairs of homologous chromosomes are fully aligned as they perform the essential meiotic task of crossover formation. A typical 2D visualization of a 3D microscopy dataset is the maximum intensity projection over the Z-axis as shown in Fig. 2A. This projection loses detail in depth, especially with data such as these nuclei, where chromosomes are radially oriented along the nuclear envelope, avoiding a large central nucleolus. Because we assume that radial organization of the signal explains most of the variation, we map the data to a sphere. This mapping is achieved by first rescaling the data to a cube of 80 pixels per side (Fig. 2B). We subsequently cast rays from the center of the cube, taking the mean intensity of the pixels along the ray. This transformation yields a dataset of the average data in spherical coordinates, the spherical projection. To compare different objects, we also normalize the spherical projection map such that the total variance is 1 and the mean is 0 (Fig. 2C).

**Figure 2.**
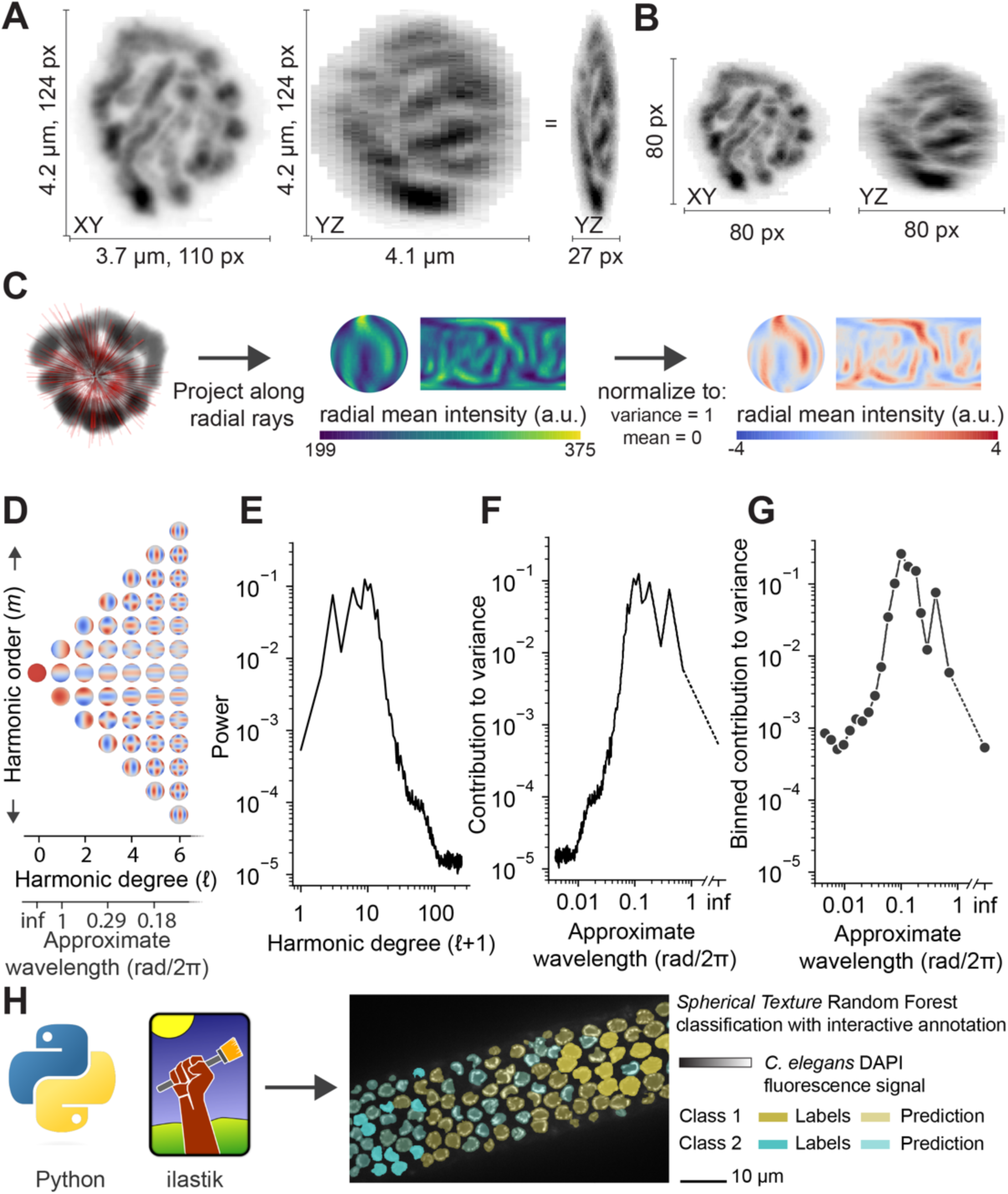
Spherical Texture method design. **A)** A *C. elegans* meiotic nucleus in the pachytene stage, stained with DAPI, shown as maximum intensity projections over Z (left) and X, with the YZ view rescaled isotropically (center) and square pixels (right) about the XY view. **B)** Data from A rescaled to 80×80×80 pixels in XY (left) and YZ (right) views **C)** A graphic showing the mean intensity projection to spherical space, showing first a subset of the radial rays (left, red lines) used to generate the mean-intensity spherical projection as spherical data and as planar map projection (center). The mean intensities are normalized to mean=0 and variance=1 (right). **D)** Projections of the spherical harmonics basis functions on the sphere of the first 7 spherical harmonic degrees. **E)** The spherical harmonics power spectrum of the spherical projection in C shows a distinct peak around approx. the 10th harmonic degree. **F**) Rescaling the harmonic degrees to approximate wavelength yields a spherical harmonics spectrum, which shows a corresponding peak in the contribution to variance around a wavelength of approx. 0.1 rad/2π **G)** The standard output of the *Spherical Texture* method corresponds to the binned spectrum shown in F. **H)** The *Spherical Texture* extraction is implemented as a Python package and directly in *ilastik*, allowing for its adoption into the Object Classification workflow. In this workflow, users can interactively train a Random Forest machine learning classifier. Shown here is a part of a *C. elegans* gonad with segmented nuclei, where some nuclei were labeled as Class 1 and others as Class 2 (solid colors). Based on the *Spherical Texture* of these labels, ilastik predicts the class of all other nuclei (transparent colors).

The spherical projection represents a meaningful dimension reduction while keeping the variation that defines the radial signal. To extract texture information from the spherical projection, we apply a Spherical Harmonics (SH) decomposition, a transformation to frequency space that is analogous to a Fourier decomposition. We decompose the spherical projection into a sum of waves (the spherical harmonics basis functions). These waves are a combination of relative scale (harmonic degree, ℓ) in different conformations (defined by harmonic order, *m*) (Fig. 2D) up to the scale of 1-pixel differences (ℓ = 251). By integrating over all harmonic orders of the signal, we get a power spectrum with a single rotationally invariant value for each degree (Fig. 2E). By normalizing to a mean of 0, the power corresponds to the variance as a function of ℓ (Wieczorek & Meschede, 2018). Therefore, the power spectrum can be reinterpreted as a measure of variance versus the approximate wavelength of each harmonic degree (Fig. 2F). Furthermore, through normalizing the projection to unit variance, the area under the curve of the *Spherical Texture* output equals 1. The method is also illustrated in video SV1.

For accessibility, we implemented the technique as a plugin for *ilastik* (Berg et al., 2019), allowing users to quickly select the *Spherical Texture* features for a Random Forest object classification algorithm. To reduce the number of features to a more relevant set, we subsample the spectrum to 20 values along the log2 axis in this implementation for further analysis (Fig. 2G-H). Here we bin these values by integrating, which ensures that the area under the curve remains equal to 1.

### Texture Extraction

To test the ability of the *Spherical Texture* technique to extract textures, we used synthetic data generated with Perlin 3D noise (Fig. 3A). This synthetic data allows us to create 3D patterns at varying spatial scales. Assessing these test patterns with the *Spherical Texture* method yields a quantification that shows how much a certain spatial scale contributes to the variance. Thus, we expect the fine patterns to have more power at small wavelengths, while the coarser patterns have the most power at larger wavelengths. Indeed, as the synthetic data gets coarser, the spectra of the *Spherical Texture* method shift from shorter wavelengths (black) towards longer wavelengths (light gray) (Fig. 3A).

**Figure 3.**
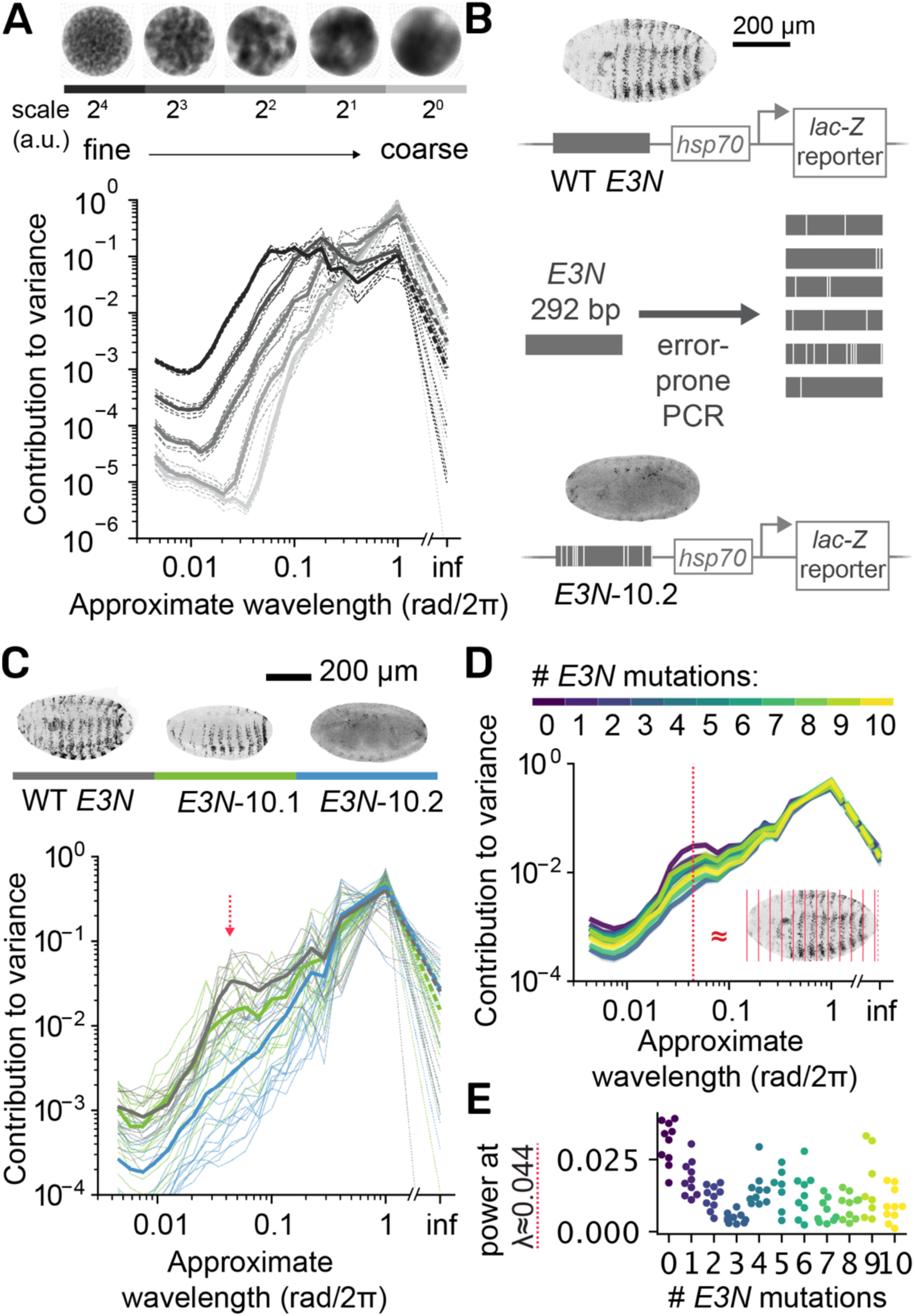
The *Spherical Textures* reflect the coarseness of 3D data and can be applied to quantify patterning in *D. melanogaster* embryos. **A)** *Spherical Textures* of synthetic 3D Perlin noise spheres. Coarser data corresponds to more variance at large wavelengths. **B)** Graphic showing the design of the mutant *E3N* enhancer screen and genetic setup. Wild-type *E3N* drives the expression of a lac-Z reporter in a striped pattern in the *D. melanogaster* embryo. By introducing mutations in the enhancer via error-prone PCR, the effect of many variants on the activity of the *E3N* enhancer can be tested by screening for changes in this pattern. C) *Spherical Texture* responses of embryos of three genotypes of the assay in B. The WT embryo (n=13) shows a distinctive average profile with a peak at λ ≈ 0.044 (red arrow), that is lost in the *E3N*-10.2 (n=17). The *E3N*-10.1 (n=18), with impaired patterning, shows an intermediate profile. **D)** Average profiles of all genotypes in the screen, clustered by number of mutations. The red dashed line is the characteristic WT wavelength, with a plane wave at the same wavelength (λ = 0.044 /object length) shown as a simplified interpretation of the wavelength. This plane wave corresponds to the distance between the stripes (inset, red stripes). **E)** The effect of different E3N enhancer variants on the gene expression pattern is described by taking the average power at λ ≈ 0.044 rad/2π for all genotypes. Separate dots are separate experiments for WT, and separate genotypes for mutants.

We next tested the ability of the *Spherical Texture* method to distinguish morphological differences in 3D microscopy images. In *D. melanogaster*, the minimal *E3N* enhancer drives *shavenbaby* (*svb*) expression in the ventral stripes of the embryo at developmental stage 15. To dissect the regulatory activity encoded in this enhancer, Fuqua et al. (2020) created a transgenic *D. melanogaster* library harboring random mutants of the *E3N* enhancer. The mutants were generated via error-prone PCR, and their activity is actualized by a downstream promoter (*hsp70*) and reporter gene (*lacZ*) (Fig. 3B). To further characterize this mutational library, a subset of 91 lines ranging from 1-10 mutations were imaged using fluorescent antibodies and confocal microscopy to study the patterns in more detail (Galupa et al., 2023). However, the analysis of high-throughput data requires an accurate and automated assessment of pattern formation. For this, the *Spherical Texture* can serve as a reliable metric. When applied to both a wild-type *E3N* reporter and two unique variants of *E3N* each harboring 10 point mutations (10.1 and 10.2), the *Spherical Texture* method distinctly differentiates between the mutants and the WT *E3N* control: the WT shows a characteristic profile, with a peak in variance at a wavelength λ ≈ 0.044 rad/2π. This peak is diminished in *E3N*-10.1 embryos which showcase less defined stripes, and it is virtually absent in *E3N*-10.2 embryos, which lost all stripe formation (Fig. 3C). We can thus effectively analyze the complete high-throughput screening data and assess the degree of pattern formation in 91 different lines (Fig. 3D-E). Our analysis reveals an abrupt decline in pattern formation fidelity from the WT strain to any of the mutated strains. However, the introduction of more than three mutations does not reveal a discernible trend. This finding suggests that the exact number of mutations (up to 10) does not define the regulatory capacity of this minimal enhancer, and some mutations may rescue other mutations in an epistatic manner.

### Classification of meiotic nuclei

To showcase a very different type of biological data, we turn to *C. elegans* germline nuclei. While the *D. melanogaster* embryos are large (500 µm diameter) and the pattern of the *E3N* enhancer is clearly defined, the *C. elegans* germline nuclei are very small (2-5 µm diameter) and lack a defined pattern. However, as the *Spherical Texture* is agnostic to the original size of the object but quantifies the scale of the morphology, we hypothesized that this method should also distinguish morphological differences in *C. elegans* germline nuclei.

*C. elegans* germline nuclei are typically categorized into three morphological stages: Proliferative zone nuclei are relatively small with chromatin distributed across the nucleus. The proliferative nuclei are mitotically dividing stem cells, generating meiotic progenitor cells. These nuclei undergo significant remodeling as they enter meiotic prophase I: the chromosomes are partially condensed and polarized within relatively small “Transition Zone” nuclei, resulting in a crescent-shaped and dense distribution of DNA. This morphological stage is indicative of the meiotic stages involving homologous chromosome pairing and synapsis. After completion of synapsis, nuclei enter the pachytene stage which is characterized by larger nuclei and separated chromosome strands representing partially condensed and synapsed homologous chromosome pairs (Hillers et al., 2017). If we apply the *Spherical Texture* to these nuclei, we find that, indeed, the *Spherical Texture* spectra represent the differences between these three morphological classes (Fig. 4A). Notably, the transition zone nuclei, where chromosomes form a large crescent-shaped structure, exhibit significantly increased variance at λ = 1 rad/2π, which implies the chromatin is organized into a half-moon-like organization. This matches the canonical description of crescent-shaped DNA morphology. For pachytene nuclei, a local peak in the spectrum is observed around λ = 0.1 rad/2π, which we infer to reflect a typical distance of separation between chromosomes in the nucleus (Fig. 4A). Thus, the *Spherical Texture* method accurately describes differences in the nuclear morphology of nuclei in the distal germline of *C. elegans*.

**Figure 4.**
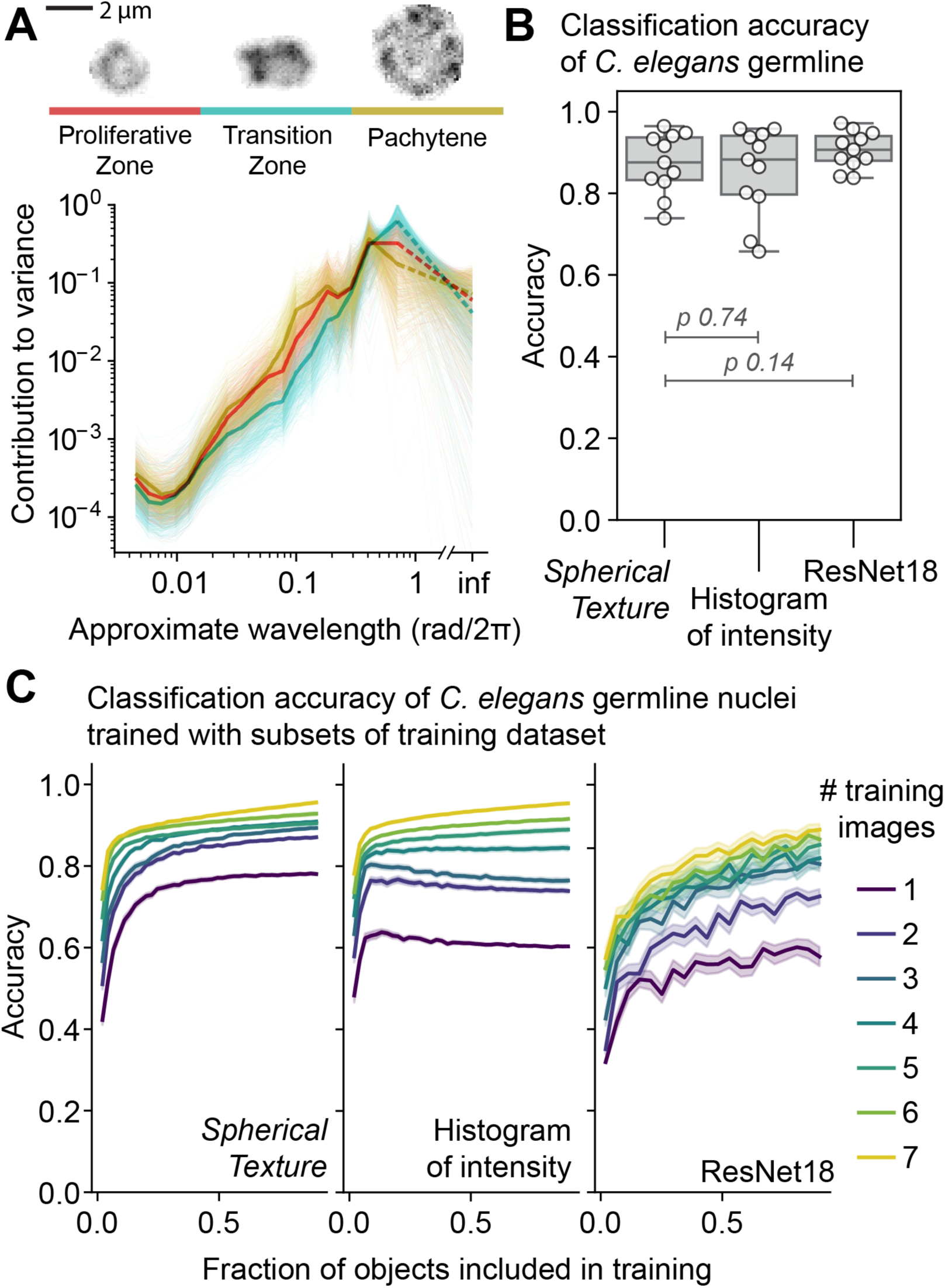
*Spherical Textures* and machine-learning classifications of *C. elegans* germline nuclei. **A)** *Spherical Texture* spectra for manually classified wild-type *C. elegans* germline nuclei show characteristic differences for each class. **B)** Classification accuracies of machine learning models classifying all annotated nuclei in one test image, trained on all annotated nuclei from 10 images. The *Spherical Texture* model is a Random Forest with the *Spherical Texture* and size in pixels as features. The *Histogram of intensity* model is a Random Forest with a 64-valued normalized histogram of intensity values and size in pixels as features. The ResNet18 is a 3D CNN with unscaled 0-padded normalized nuclei at original scale as input. The models behave similarly, but the ResNet slightly outperforms the Random Forest models as expected. Stated *p-*values are from a Wilcoxon one-sided paired test, testing for accuracy greater than *Spherical Texture*. **C)** The classification accuracy increases with increasing amount of training data for the three models. The color denotes the number of images used for training, and the x-axis represents the fraction of nuclei from each these images. Only the *Spherical Texture* trains monotonically and quickly, while the Histogram of intensity overfits with few images, and the ResNet requires a large amount of training data to reach high accuracy.

We then utilized the differences identified in the *Spherical Texture* spectra to classify the different stages of nuclei within the *C. elegans* distal germline. To achieve this, we used a machine-learning approach by training a Random Forest classifier. Random Forest classifiers are simple and minimal to set up, and implemented in available software such as *ilastik*, providing user-friendly interactive image classification and analysis (Berg et al., 2019). We included the *Spherical Texture* spectrum (Fig. 2E) and the original size of the nuclei as features. We compared this *Spherical Texture* classification to a Random Forest using the *ilastik* histogram of intensities and nucleus size as a feature set, or a more complex convolutional neural network model, a 3D ResNet18 (He et al., 2015), that learns a feature set from the 3D segmented nuclei. After training on 10 different annotated images containing over 1400 annotations, we found that all models had similar levels of accuracy. However, the ResNet was the most consistent among them (Fig. 4B).

In bioimaging, the amount of training data is often limiting, as experimental techniques, the time required for annotation, and the inconsistency in experimental conditions all hinder the generation of comprehensive and consistent training datasets. Consequently, the efficiency of model training becomes a critical consideration. We systematically shuffled and subsampled our training set by the number of images and included objects, generating smaller subsets of our cross-validation dataset. By creating these smaller subsets, we were able to investigate how well the models learn to classify germline nuclei when training data is limited (Fig. 4C, S1). This analysis reveals that the *Spherical Texture* model exhibits fast and consistent training that improves monotonically with increasing training data size. The *Histogram of intensity* model also trains rapidly, but the accuracy declines as more data is added from a limited number of images. This decline is likely due to the highly sample-specific variations in fluorescence intensities which can lead to overfitting when only training on a small set of images. In contrast, the ResNet model, while accurate when trained on the full dataset, was far less accurate when provided with less training data, which is consistent with evaluations of 2D ResNet models (Brigato & Iocchi, 2020).

We can now leverage the *Spherical Texture* model for germline classification to assess meiotic staging in the *C. elegans* gonad. This is feasible due to the temporo-spatial organization of the *C. elegans* gonad, wherein nuclei progress through the gonad while undergoing meiosis. Thus, the nuclei are separated into phenotypic zones (Hillers et al., 2017). Traditionally, the manual annotation of these zones relies on marking the transition points where most nuclei change morphology (Fig. 5A). However, this approach becomes challenging, and at times biased, especially when genetic defects give rise to gradual transitions. We therefore use the *Spherical Texture* model, consisting of a Random Forest trained using the *Spherical Texture* spectrum and nucleus size as features to predict the stages of nuclei in germlines of wild-type animals (Fig. 5A, higher resolution in S2). Indeed, we find that the *Spherical Texture* classifications of individual nuclei of wild-type germlines mostly match the zones expected from the overall germline organization: nuclei in the distal (here shown left) part of the gonad are classified as “proliferative zone” nuclei, moving proximally to first transition zone and then pachytene nuclei. However, as the nuclei exit the transition zone and shift to early pachytene, some early pachytene nuclei are misclassified as “proliferative zone” nuclei. This finding suggests that the *Spherical Texture* model not only identifies the three canonical zones but also detects morphological differences between nuclei in early and mid/late pachytene, respectively.

**Figure 5.**
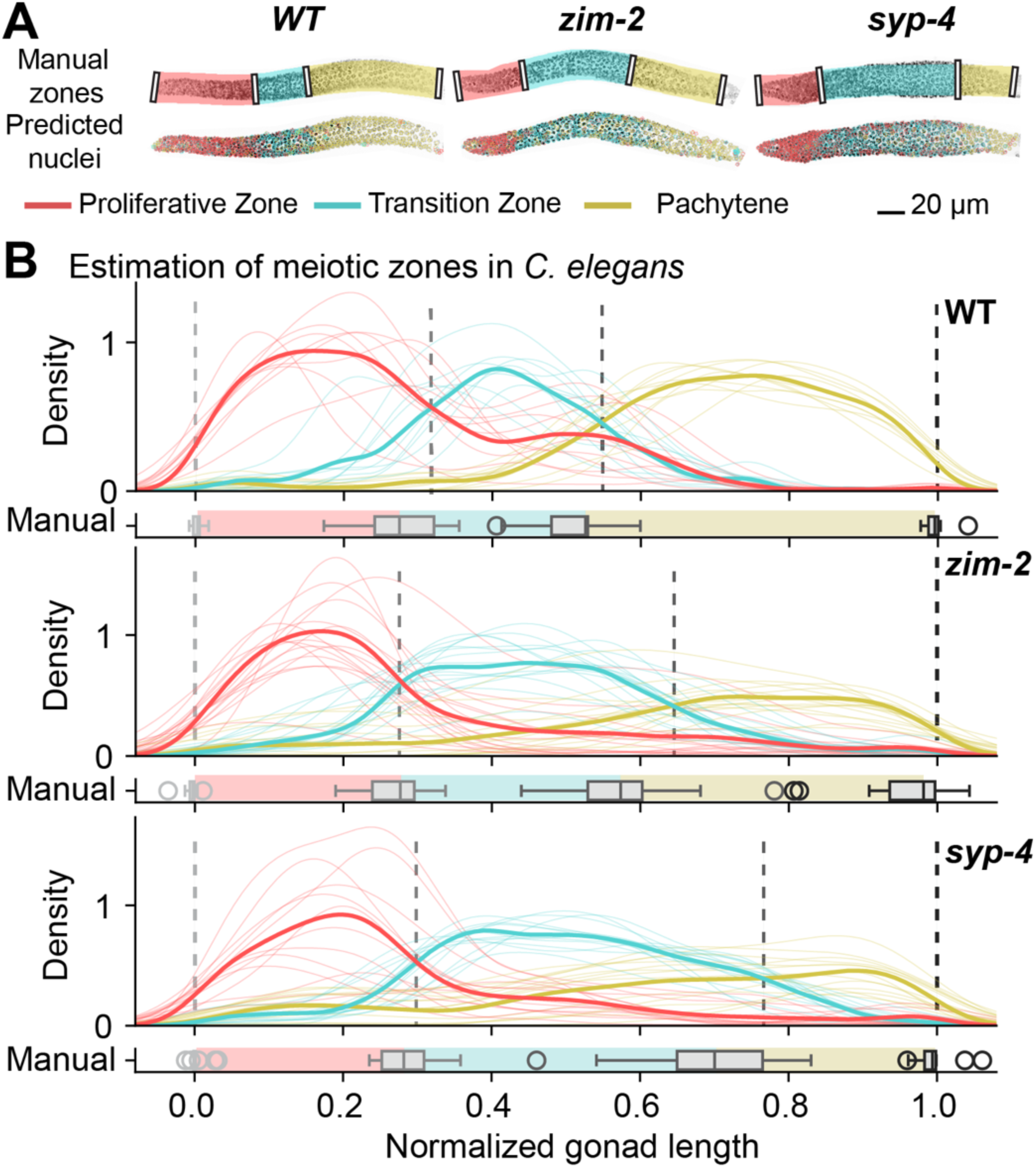
Automatic classification of germline nuclei provides quantifications of meiotic progression. **A)** Representative images of *C. elegans* gonads of three genotypes (Wild-type, *zim-2, syp-4*) with manually annotated zones, and automatic classifications of nuclear morphology using the *Spherical Texture* Random Forest model per nucleus. Higher resolution images are available in S2. **B)** Bulk analysis of *Spherical Texture* annotations in gonads reveals average zone sizes along the linearized gonad. The relative density distribution of nuclei per morphological classification is plotted along the gonad central spline. The point where the means of the zones cross are annotated (dashed lines) to compare against the manual annotations of these transition points.

Due to the temporo-spatial organization of the *C. elegans* germline, the length of individual zones corresponds to the time individual nuclei spend within each stage (Hillers et al., 2017). Therefore, the length of the transition zone within a gonad is a reliable metric to determine the timely completion of homologous chromosome pairing and synapsis that take place in this zone. In animals with mutations in genes involved in pairing or synapsis, the transition zone length is altered. For instance, in *zim-2* mutant animals pairing, and consequently synapsis, of a single chromosome, namely chromosome V, is eliminated (Phillips & Dernburg, 2006), while in *syp-4* mutant animals, synapsis is completely abolished (Smolikov et al., 2009). Despite being trained solely on nuclei of wild-type germlines, the *Spherical Texture* method, predicts elongated transition zones for both *zim-2* and *syp-4* animals (Fig. 5A).

With the *Spherical Texture*-based model, we can analyze meiotic progression across many different animals by automatically classifying all nuclei across many gonads allowing for automatic quantification of transition zone length (Fig. 5B). Notably, we observe robust progression through the three zones which matches manual annotations. The machine-learning-based prediction pinpoints not only the most probable position of the transition between zones but also illustrates the steepness of this transition. In wild-type animals, shifts between zones occur rapidly, while the progression from the transition zone to pachytene is more gradual in both mutant animals. As a result, the *Spherical Texture* method predicts an even longer transition zone for both mutants compared to our manual annotations.

Utilizing the *Spherical Texture* predictions along the length of the gonad offers a clear, highly informative, and easily interpretable representation of meiotic zones in *C. elegans*. This method allows for a consistent analysis of large datasets in a streamlined manner. Moreover, due to the model’s fast training speed, the *Spherical Texture* model can easily be adapted to other imaging modalities or experimental conditions.

### 2D texture and polarization quantification

Despite the popularity of 3D imaging, 2D imaging remains a prevalent and valuable tool for addressing various biological questions. Similar to 3D, patterning and the distribution of intensity across objects remain central features also in 2D. To analyze textures in 2D data with the *Spherical Texture* method, we can project the intensities within the convex region of an object to a circle instead of a sphere decomposing with a 1D Fourier transform (Fig. 6A). This process results in a power spectrum of the projection, which yields a quantification depicting the contribution to variance per wavelength, where the area under the curve equals 1 when normalized to mean 0 and variance 1. When applied to 2D Perlin noise patterns corresponding to the middle slices of the 3D synthetic data depicted in Fig. 3A, a similar distribution of 2D power spectra emerges, mirroring what we observed for 3D spherical data: the peak of the power spectrum moves from short wavelengths to longer wavelengths as the data becomes coarser (Fig. 6B). Therefore, our *Spherical Texture* method efficiently quantifies textures not only in 3D but also 2D.

**Figure 6.**
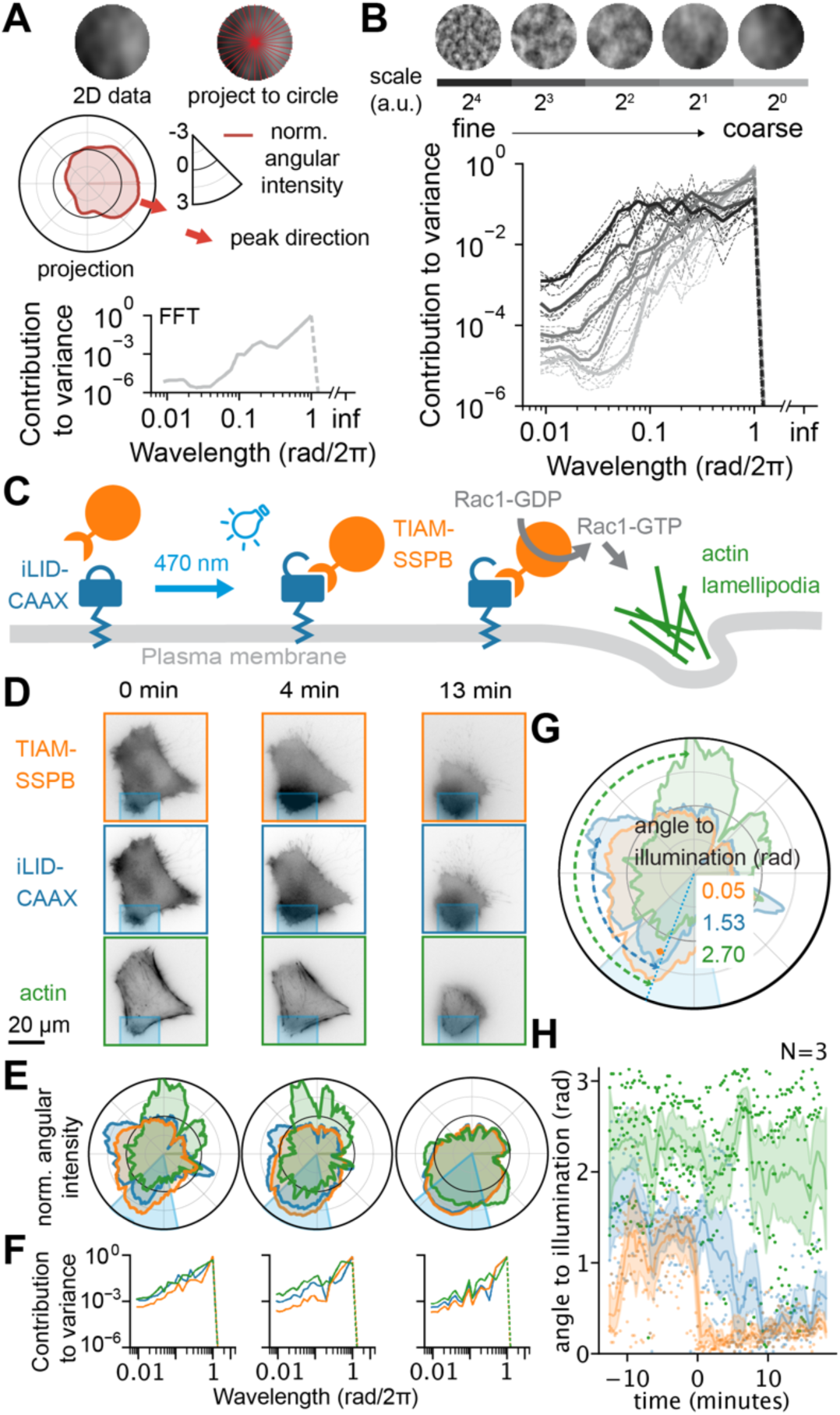
Texture and polarization extraction of 2D data. **A)** Explanation graphic of the *Spherical Texture* for 2D data (top left) – the data is sampled per angle (top right) and projected as mean intensity to a 1D circle (S^1^ space, middle), and normalized to unit variance, mean 0. The contribution to variance per wavelength is then calculated through the Fourier power spectrum (bottom). The ‘peak direction’ (red arrow) denotes the angle of the maximum value in the projection. **B)** 2D *Spherical Texture* spectra for synthetic Perlin noise circles. The finer-detailed spectra have more power at short wavelengths, while the coarse spectra only have power at long wavelengths. **C)** Graphic depicting the optogenetic system for Rac1 activation (de Beco, 2018). The membrane anchor CAAX (blue) is tagged with the photosensitive domain iLiD, which – upon activation with 470 nm light – recruits TIAM-SSPB (orange). TIAM acts as a guanine nucleotide exchange factor for Rac1 (gray). The Rac1-GTP induces the formation of actin-rich lamellipodia (green). **D)** Fluorescence images with **E)** projections (same axis as A) and **F)** spectra of an illuminated cell. The illuminated region is shown in cyan. Three different time points are shown from left to right. As time post-illumination progresses, both the TIAM (orange) and CAAX (blue) align to the illumination. By contrast, actin (green) switches from a broader distribution to an almost bipolar distribution with peaks at the site of illumination, and directly opposite. These changes are also reflected in the spectra where TIAM and CAAX gain power at 1 rad/2π and actin at 0.5 rad/2π. **G)** The graphic illustrates the measurement of the angle to illumination which denotes the shortest angle between the peak direction of individual channels and the peak of the illumination. Therefore, the maximum angle to illumination is π. **H)** Angle to illumination for all channels of three cells over time (illumination at 0 min). TIAM aligns immediately to the illumination angle, while CAAX aligns slower. Actin splits into two populations: one aligning to the illumination, and one that aligns directly opposite the illumination reflecting its bipolar distribution.

An application for employing this method in 2D is in quantifying actin dynamics during cell polarization. In such assays, cells polarize forming a distinct ‘leading edge’ characterized by actin-rich lamellipodia oriented towards the direction of movement. To precisely define the position of the leading edge, we utilize an optogenetic approach. Here, a photosensitive tag on the membrane anchor CAAX recruits TIAM, a Rac1-specific guanine nucleotide exchange factor, upon light stimulation. With this setup, Rac1 can be activated at a specific site, thereby inducing leading edge formation at that exact site (Fig. 6C, Video SV2) (de Beco et al., 2018). We aim to quantify leading edge formation in this system, by using the *Spherical Texture* method on the live-cell actin probe SiR-actin to visualize actin dynamics and polarization together with CAAX and TIAM.

In unstimulated cells, both TIAM (orange) and CAAX (blue) are randomly distributed across the cell (Fig. 6D, 0 min). This random intensity distribution shows up as random fluctuations in circular intensity projections (Fig. 6E, 0 min), leading to an unstructured variance spectrum in the *Spherical Texture* quantification (Fig. 6F, 0 min,). Upon stimulation at 0 min, TIAM quickly accumulates at the activation site which is reflected by a higher contribution of larger wavelength to the variance of the signal reflecting the coarsening in the distribution of the TIAM signal (Fig. 6D-F, at 4 and 13 min). Interestingly, the CAAX signal also intensifies at the activation site and almost completely overlaps with the TIAM signal. This observation is intriguing because CAAX is not specifically recruited by light stimulation. We infer that this apparent accumulation is a consequence of membrane ruffling and lamellipodia formation. In the circular projection at minute 4, the signals of TIAM and CAAX overlap, while we observe that the actin intensity is only slightly increased at the activated site and is, instead, concentrated at the rear of the cell as it retracts. However, at minute 13 we observe a clear accumulation of actin at the activation site in the circular projection, indicating the polarization of the cell.

To analyze this further, we measure the angle between the illumination and the polarization direction of the circular projections to evaluate the alignment of TIAM, CAAX, or actin relative to the illumination (Fig. 6E). Assessing these angles across three cells over time reveals that TIAM aligns with the illumination almost immediately (time 0), and CAAX aligns within a few minutes until it is fully aligned about 10 minutes post-illumination (Fig. 6F). By contrast, the angle between illumination and peak actin intensity remains large throughout the imaging time. However, upon closer examination, we find a bimodal distribution of the angles with some peak intensities located at the activation site while most cluster around π, indicating that actin accumulates both at the activation site and at the rear end opposite the activation site, as previously observed in the circular projections. A modest accumulation of actin at the site of illumination is consistent with induced Rac1-mediated branched actin polymerization and lamellipodia formation. We infer that the increase in actin seen at the opposite side of the cell is consistent with a restructuring of the cell membrane and morphological changes, as a migratory rear edge is formed, where the intensity subsequently diminishes over time.

## Discussion

Here, we presented a texture extraction method designed for the quantification and classification of objects in microscopy images. This method efficiently extracts texture resolution from both 3D and 2D objects, operating under the assumption that many biological objects are largely convex and can be described as angular variations from their centroid. Our study showcases the effectiveness of this technique for diverse applications ranging from pattern recognition in 3D images of *D. melanogaster* embryos to the quantification of 3D nuclear morphology in *C. elegans* gonads. Furthermore, our texture analysis approach extends to 2D scenarios such as real-time images of migratory cells. When coupled with signal analysis and peak finding in circular projections, it provides a measure for cell polarization and migration direction.

The *Spherical Texture* method yields a reliable metric to quantify pattern formation in gene expression driven by the E3N enhancer in the *Drosophila melanogaster* embryo, which features a clear and predefined biological pattern. The rotationally invariant signal produced by the *Spherical Texture* method allows for robust and consistent quantification that is independent of the orientation of the input image of the fly embryo. This independence of sample mounting on the quantification result makes the *Spherical Texture* method ideal for analysis of large-scale screens acquired by automated imaging.

On the other hand, nuclei in the distal *C. elegans* germline lack a predefined pattern but exhibit general differences in their morphology that is a consequence of differences in DNA condensation and nuclear organization. The *Spherical Texture* method extracts features based on their scale which allows for the robust classification of nuclei based on their morphology - a task that was previously only achievable by manual annotation. Therefore, the *Spherical Texture* method can be applied to both structured patterns such as patterning during *Drosophila* embryogenesis, and unpatterned data such as nuclear morphologies in the *C. elegans* germline highlighting its versatility.

To utilize the texture information obtained by our *Spherical texture* method for object classification, we integrated this tool into the easy-to-use interactive learning and segmentation software *ilastik* (Berg et al., 2019). Employing Random Forests using features derived from the *Spherical Texture* method to classify *C. elegans* nuclei demonstrated consistent performance and outperformed more complex CNNs in scenarios with sparse training data. Even with a dataset of over 1400 annotations, the CNN-based model only marginally surpasses the consistency of the Random Forest model. This is particularly relevant since microscopy datasets are highly specific to the lab, microscope, and experiment, necessitating frequent (re)training. Additionally, the challenges of implementing a 3D CNN architecture require more expert knowledge and contrast with the ease of using a Random Forest, especially within software like *ilastik*.

The incorporation of the *Spherical Texture* method into *ilastik* also facilitates the seamless combination with other object quantification features. At the same time, the *Spherical Texture* method can be combined with other signal analysis techniques applied to the radial projections. We illustrate this approach in the actin leading edge quantification. Peak finding algorithms applied to the *Spherical Texture* output of the fluorescent image of a migratory cell after circular projection allow for efficient measurements of cell polarization, providing information on both the direction and relative intensity of the leading edge. This peak-finding feature in circular or spherical projections can be added to *ilastik* as a custom feature and can be used for both 2D and 3D data.

We therefore anticipate that our *Spherical Texture* method makes texture extraction easily accessible to users, and allows for its application to diverse datasets.

## Methods

### Spherical Texture implementation

The *Spherical Texture* quantification requires either 3D z-stack or 2D image data and the corresponding segmentation masks where the centroid is inside the mask.

For each object in the segmentation mask, the image data is scaled bilinearly to 80×80×80 (or 80×80 in 2D) and masked out with the provided segmentation mask. Spherical rays are taken from the centroid to angles fitting a Gauss-Legendre Quadrature (Wieczorek & Meschede, 2018). This process yields a spherical projection of 251 by 512 rays. To obtain the value of a pixel in the spherical mean intensity projection, pixel intensities are averaged along each ray until it leaves the segmentation mask. Subsequently, the spherical projection map is normalized to achieve a mean of 0 and a variance of 1 using the formula 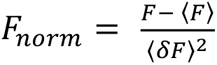, where F represents the angular mean intensity projection. The normalized signal is then decomposed into geodesy 4-pi normalized spherical harmonics using the SHTOOLS 4.10.3 (Wieczorek & Meschede, 2018) implementation of the Holmes and Featherstone algorithm (Holmes & Featherstone, 2002). Spectra are binned along a log_2_ scale to produce 20 unique output values. Binning is performed through local integration to retain the area under the curve. Given that the mean of the signal is 0, the resulting spectrum can be interpreted as variance as a function of harmonic degree ℓ (Wieczorek & Meschede, 2018).

To map the harmonic degree ℓ to the approximate cartesian wavelength λ, we use the Jeans relation 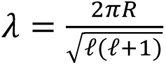 for the unit sphere with radius R = 1 (Wieczorek & Meschede, 2018). This relation does not hold well for lower values of ℓ. To address this limitation, we set the cartesian wavelength λ = 1 for ℓ = 1, where the basis functions exhibit only one peak and one valley across the sphere. For ℓ values greater than 1, we rely on the Jeans relation for simplicity.

For two-dimensional data, we cast the rays only along the equator, resulting in a circular line comprising 251 values. This line is then decomposed using the discrete Fourier Transform implementation available in scipy (Virtanen et al., 2020).

Polarization directions are calculated from the angle of the maximum value in the projection. Depending on data, it might be effective to first bandpass the signal.

### Integration into ilastik and code availability

The *Spherical Texture* method is in the latest version of *ilastik*, starting at 1.4.1b19 (Berg et al., 2019). The *ilastik* implementation of *Spherical Textures* is integrated into the *ilastik* Object Classification workflow, where users provide a 2- or 3D image and segmentation mask for which different features can be extracted. The *Spherical Texture* spectrum and peak extraction are both easily selectable by checking the appropriate checkboxes. Within the software, one can then interactively train a Random Forest classifier to classify phenotypes based on the extracted features. The extracted features, including the *Spherical Textures*, can also be exported separately and used in subsequent custom analyses.

The code is implemented in Python and accelerated with numba (Lam et al., 2015) with parallel computation of multiple objects. For users who want direct access to the code in Python, outside of the *ilastik* implementation, a Python package is installable through pip and conda as described on https://github.com/KoehlerLab/SphericalTexture.

### Synthetic data generation

3D Perlin noise (Perlin, 1985) was generated in 128×128×128 pixel grids using the perlin-numpy python implementation (Vigier, 2018/2024). The noise scale parameter is the relative ‘periods of noise’ generated along each axis across the 128 grid. By design, Perlin noise periods are relatively arbitrary and do not decompose into clean waves. To obtain spherical synthetic data, we provided a central 80×80×80 sphere as a mask. For 2D synthetic data, only the middle plane was used.

### Fly strains, reporter gene expression staining and imaging

As previously described in Galupa et al. (2023), a subset of 91 lines of the original 749 variants, ranging from 1-10 mutations, of the mutant library generated by Fuqua et al. (2020) was used for this analysis. These transgenic *Drosophila melanogaster* lines were based on *attP2* (Bloomington Stock Number: 5905). Fly rearing, embryo collection and fixation, and immunofluorescence was performed as described before (Fuqua et al., 2020; Galupa et al., 2023). Z-stacks of every embryo were acquired using a confocal Zeiss LSM 880 microscope at 0.593×0.593×1.40 µm pixel size using a 20x 0.8 NA air plan-apochromatic objective.

Masks were created in the 2D maximum intensity projections of the data using the ‘cyto’ pretrained cellpose model with a target diameter of 600 pixels, corresponding to 356 µm (Pachitariu & Stringer, 2022; Stringer et al., 2021). These masks were then extended through the Z dimension.

### Imaging of C. elegans germlines

To visualize *C. elegans* germline nuclei, young adult N2, CV87 [*syp-4*(*tm2713*)], or CA258 [*zim-2*(*tm574*)] animals were dissected 24 hours post-L4 and stained with DAPI (Sigma-Aldrich, D9542) as previously described (Köhler et al., 2017; Phillips et al., 2009). Dissected gonads were mounted in ProLong Glass antifade mounting medium (Invitrogen, P36984). Images were acquired on an Olympus spinning disk confocal IXplore SpinSR system using a 60X 1.4 NA oil plan-apochromatic objective. High-resolution images for Fig. 1B and Fig. 2 were acquired with a SoRa disk at a 0.034×0.034×0.16 µm pixel size, while all other images of germline nuclei were acquired with a 50 µm disk and a 0.108×0.108×0.2 µm pixel size.

Nuclei were segmented using a customized cellpose model (https://github.com/KoehlerLab/Cellpose_germlineNuclei/blob/main/Cellpose_germlineNucl ei/cellpose_germlineNuclei_KoehlerLab) as previously described (Piñeiro López et al., 2023). Regions containing distal germline nuclei from proliferative zone until the end of pachytene were manually annotated in Fiji (Schindelin et al., 2012), and only masks within this region were used in all downstream analyses.

### Gonad linearization

Gonads were linearized by fitting a cubic spline to a LOWESS fit of the positions of segmented objects larger than 10 pixels in a manually annotated region of the gonad, excluding somatic cells. Nuclei position along gonad length is defined as the point along the spline where the distance to the nucleus is minimal.

### Manual annotation of C. elegans germline nuclei

1665 nuclei were annotated in 11 gonads of WT *C. elegans* in *ilastik*, without the feedback of the *ilastik* interactive labeling to not bias the cross-validation dataset. Nuclei of unclear phenotypes or nuclei with incorrect segmentations were ignored in the annotation.

### ResNet implementation

A 3D-ResNet was constructed from the pytorch implementation of ResNet18 (He et al., 2015), by changing the 2D convolutions into 3D convolutions. To give it similar information as the Random Forests, the data sent to the ResNet were masked segmented nuclei, normalized between −1 and 1, and 0-padded to the size of the largest nucleus. Thus, the relative size of individual nuclei is retained in the image data.

For Fig. 4B, where almost the whole dataset was used as training, the models were trained for 100 epochs, with the accuracy saturating already at around 25 epochs. Therefore, all other models in 4C were only trained for 25 epochs.

### Random Forest models

To classify germline nuclei, we generated Random Forests using the default scikit-learn implementation (Pedregosa et al., 2011) with 100 estimators. We used the 20-value *Spherical Texture* output and the size in pixels of each object (total number of pixels) for the *Spherical Texture* model, or a 64-value normalized histogram as is the default in *ilastik* and the size in pixels of each object for the *Histogram of intensities* model as features.

### Photoactivation

HT1080 fibrosarcoma cells (ATCC) were cultured in DMEM supplemented with 10% FBS and 50 µg/ml penicillin/streptomycin at 37°C in 5% CO_2_. A stable cell line for optogenetic TIAM recruitment was produced using lentiviral transduction. pLenti-TIAM-tagRFP-SSPB-P2A-mVenus-iLID-CAAX was used for production (a gift from M. Coppey).

Lentivirus were produced by transfecting 10 cm dishes of HEK293T cells with 15µg pLenti construct, 10 µg psPAX2 lentivirus packaging plasmid (a gift from Didier Trono, Addgene #12260) and 5 µg lentivirus envelope plasmid (a gift from Didier Trono, Addgene #12259) with 90 µL 1 mg/mL MaxPEI. 24 and 48 hours following transfection, viral supernatant was harvested, filtered with a 0.45 µm syringe filter, and precipitated in 1X virus precipitation solution (from 5X solution: 66.6 mM PEG 6000, 410 mM NaCl, in ddH_2_O, pH 7.2). Following storage at 4°C for 24 hours, the viral supernatant was centrifuged for 30 min at 1500 x g at 4°C, and the virus pellet was resuspended in 1X PBS for long term storage at −80°C.Wild-type HT1080 cells were used as a target for lentiviral transduction. 24 h prior to transduction, HT1080 cells were seeded to a 24-well plate. On the day of transduction, the medium was refreshed with complete medium with 5 µg/mL polybrene, and 5 µL viral suspension was added. The medium was refreshed after 24 hours, and cells were selected in complete medium with 20 µg/mL blasticidin.

18 hours prior to imaging, cells were plated on 25 mm coverslips and incubated with 10 nM SiR-actin in complete medium. Epifluorescent images with photostimulation were acquired using a Nikon Ti inverted microscope equipped with a 40× (Plan Fluor, NA 1.3) oil objective, sample incubator (Tokai-Hit), ET 514-nm Laser Bandpass (49905), ET-mCherry (49008) and ET-Cy5 (49006) filter cubes (all Chroma), pco.edge cooled sCMOS camera (Excelitas) and a Polygon 400 digital mirror device (Mightex). µManager 1.4 (Edelstein et al., 2010) was used for controling the microscope, and Polyscan 2 (Mightex) was used for light patterns. Light exposure was synchronized with camera frames using camera-evoked TTL triggers. Cells were imaged with a 15 s interval and stimulated in a local region of interest with 2 mW/cm^2^ 470 nm LED (Mightex) between imaging frames.

## Data availability

All code and software are available as indicated in the Material and Methods section. Data acquired for this manuscript are available at https://doi.org/10.5281/zenodo.12745516.

## Acknowledgments

We thank Ivana Čavka for help with gonad linearization, Johannes Hugger for adapting the pytorch ResNet for 3D data, and Niccolò Banterle for discussions. We thank the high-performance computing cluster at the European Molecular Biology Laboratory (EMBL) for its support. The EMBL Advanced Light Microscopy Facility (ALMF), Zeiss, and Evident/Olympus are acknowledged for support in image acquisition. Some *C. elegans* strains were provided by the Caenorhabditis Genetics Center, which is funded by the National Institutes of Health Office of Research Infrastructure Programs (P40 OD010440).

This work was supported by: the European Molecular Biology Laboratory (SK, AK, JC), and by the Deutsche Forschungsgemeinschaft (DFG, German Research Foundation) - project number 452616889 (SK). JP, WN and LCK were supported by the Eindhoven, Wageningen Utrecht Alliance through the Centre for Living Technologies.

## Author contributions

Conceptualization: OG, SK

Formal analysis: OG

Investigation: JBP, NB, WN, SK

Data curation: OG

Software: OG Methodology: OG, SK, DK

Visualization: OG

Supervision: JC, LCK, AK, SK

Writing – original draft: OG

Writing – review & editing: OG, JBP, NB, DK, WN, TF, JC, LCK, AK, SK

## Supplemental data

**Supplemental Figure S1.**
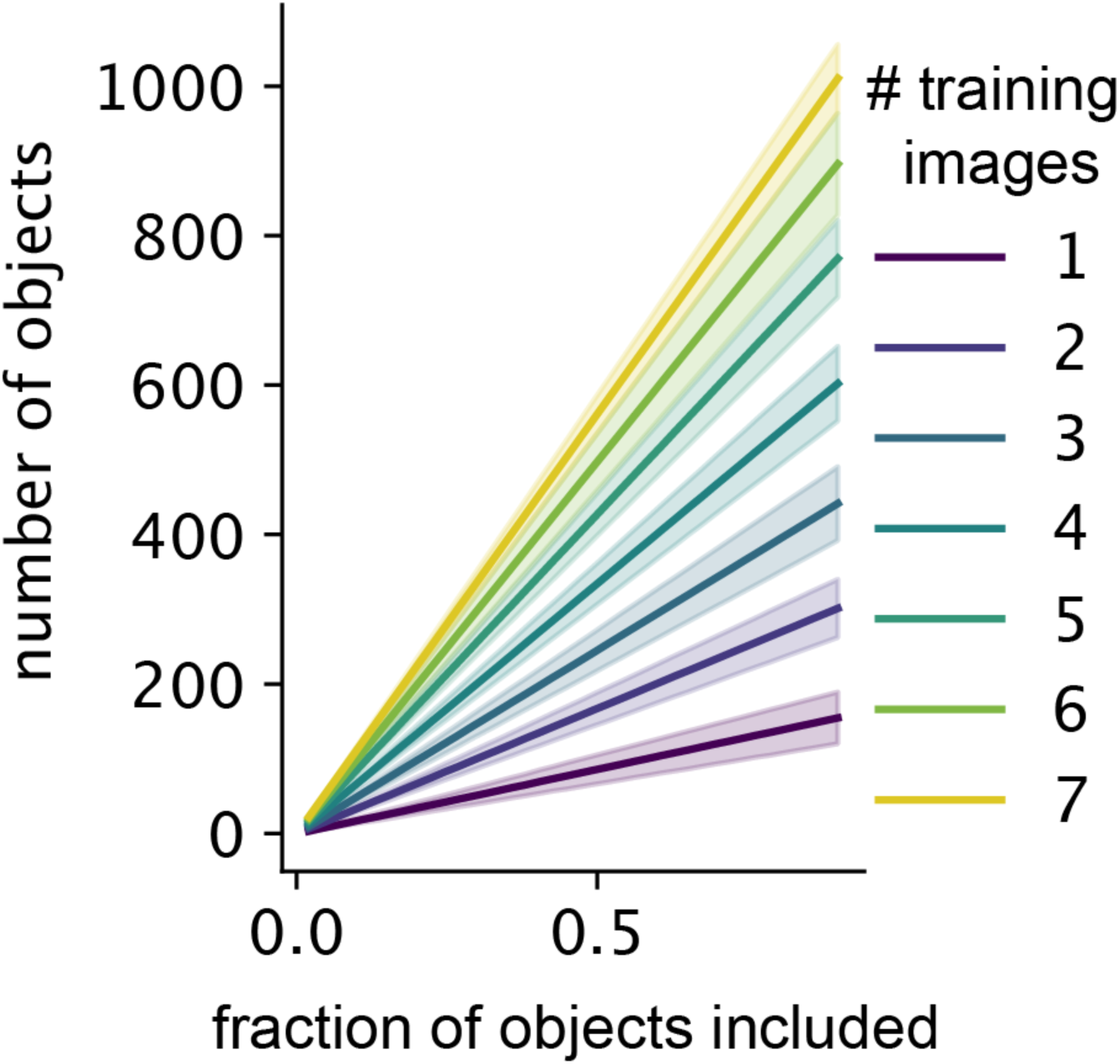
The average number of nuclei included for subsetting the *C. elegans* germline nucleus training dataset shown in Fig. 4C.

**Supplemental Figure S2.**
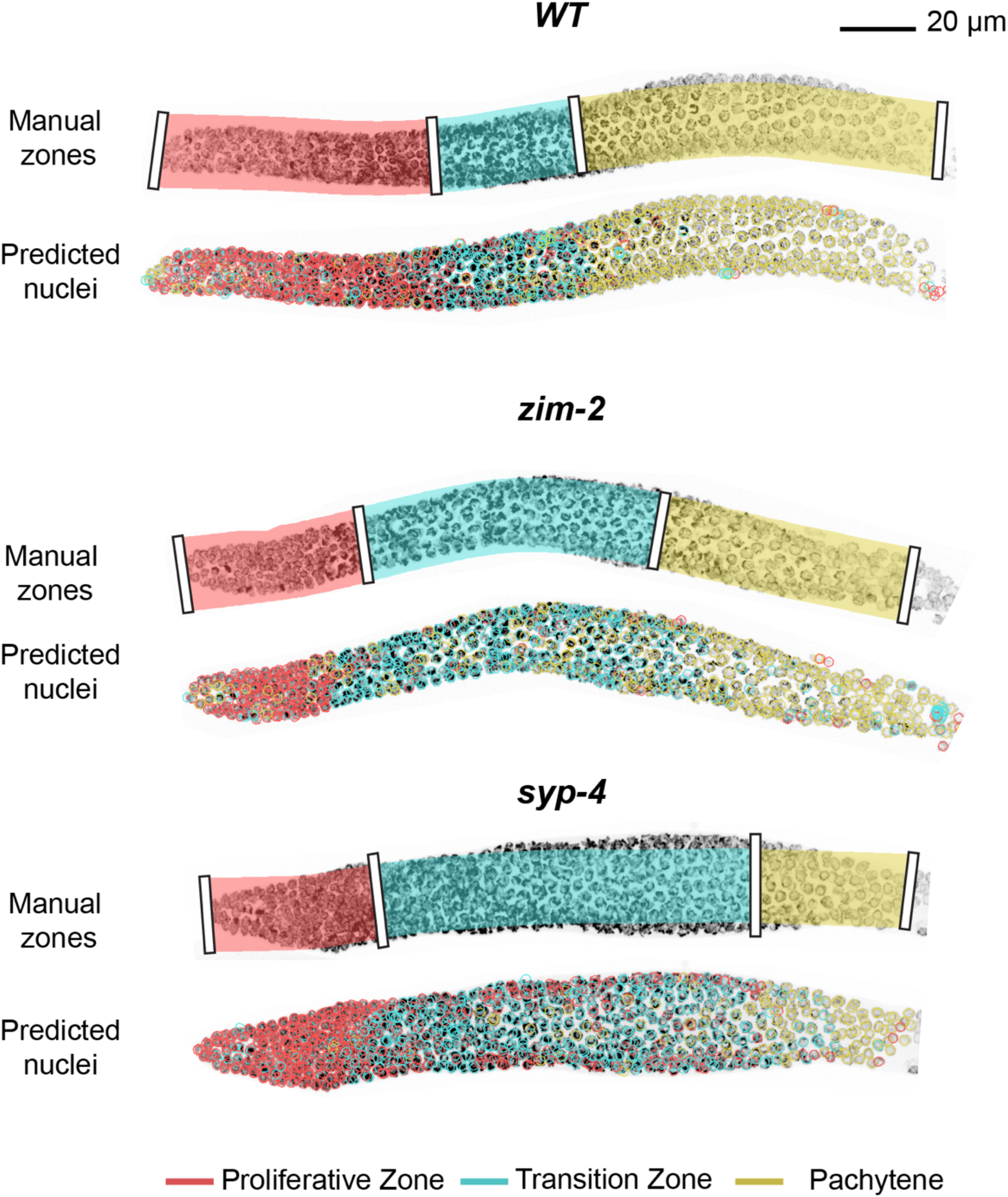
High-resolution images of *C. elegans* germlines shown in Fig. 5A for representative images from three genotypes (Wild-type, *zim-2*, *syp-4*) with manually annotated zones (top) and automatic classifications of nuclear morphology using the Spherical Texture Random Forest model per nucleus (bottom).

**Supplemental Video SV1.** Illustration of the method showing the analysis of a single *C. elegans* nucleus to the *Spherical Texture* spectrum.

**Supplemental Video SV2.** Video of photoactivation of a fibrosarcoma cell with quantification. **A)** Fluorescence video of TIAM-SSPB channel **B)** Fluorescence video of iLiD-CAAX channel **C)** Fluorescence video of SiR-actin channel **D)** Normalized angular intensity of all channels **E)** *Spherical Texture* spectrum of all channels

